# Short-Term Meditation Training Alters Brain Activity and Sympathetic Responses at Rest, but not during the meditation

**DOI:** 10.1101/2023.12.01.569603

**Authors:** Anna Rusinova, Maria Volodina, Alexei Ossadtchi

## Abstract

Numerous studies have shown that meditation has a number of positive effects on the physical and psychological well-being of practitioners. As a result, meditation has become widely practiced not only as a religious practice but also as a self-regulation technique to achieve specific measurable goals. This raises the question of how quickly physiological changes can be noticed in individuals for whom meditation is not the main focus of their lives but rather a wellbeing keeping technique. Another question is whether it is possible to observe changes occurring directly during meditation and use bio- or neuro-feedback to enhance such meditation training and achieve tangible results.

In our study, the experimental group of individuals with no previous meditation experience underwent eight weeks of training in Taoist meditation (2 sessions lasting 1 hour each week), under the guidance of a certified instructor. Participants in the control group attended offline group meetings during the same period, where they listened to audio books. All participants performed meditation testing before and after the intervention, following audio instructions. During the meditation practice, participants’ EEG, photoplethysmogram, respiratory rate, and skin conductance were recorded.

The meditation training, but not the control group activity, resulted in topically organized changes of the resting state brain activity and heart rate variability. Specifically, we observed an increase in EEG power in multiple frequency bands (delta, theta, alpha, beta) and changes in the heart rate variability indicators associated with sympathetic system activation. However, no significant changes were observed when we compared the physiological indicators during the actual meditation process performed prior and post the 8-week training. We interpret these changes as signs of increased alertness and possibly accelerated resting metabolic rate. Importantly, these changes were observed after only 16 hours of meditation training performed during the 8-week period of time. The absence of difference in the band-specific power profiles between the experimental and control groups during the process of meditation conceptually complicates the development of assistive devices aimed at “guiding” the novice meditators during the actual meditation. Our results suggest that the focus in creating such digital assistants should rather be shifted towards monitoring neurophysiological activity during the time intervals outside of the actual meditation. The apparent changes occur not only in the EEG derived parameters but are also detectable based on the markers of autonomous nervous system activity that can be readily registered with a range of wearable gadgets which renders hope for a rapid translation of our results into practical applications.

## 1. Introduction

Meditation has a long history as a self-regulation practice aimed at alleviating mental suffering and promoting enlightenment (Goenka 2012). Over time, the perception of meditation has shifted from being solely for spiritual pursuits to a means of promoting well-being and reducing stress (Gupta and Kumar 2021). Research suggests that meditation can lead to improvements in cognitive abilities such as attention, concentration, creativity, and problem-solving (Walsh and Shapiro 2006). It has also been found to contribute to changes in brain function and structure (Davanger et al. 2010), (Farb, Segal, and Anderson 2013), (Hölzel et al. 2011), (Kang et al. 2013) and autonomic activity (Kubota et al. 2001).

Meditation studies focus on two main effects: the physiological changes observed during meditation, and the long-term effects of consistent meditation practice. Whereas investigating the long-term effects of meditation is crucial for understanding the underlying mechanisms behind the reported benefits of meditation, analyzing the physiological changes during meditation is important for the development of monitoring meditation devices to ensure a smooth and harmless meditation experience (Ahani et al. 2014).

Analyzing the long-term effects of meditation can be aided by conducting studies with experienced meditators. There is evidence that experienced meditators in a resting state, unlike novices, are calmer, breathe more slowly, exhibit increased brain activation in certain neuroanatomical areas (Vestergaard-Poulsen et al. 2009), have improved emotional stability (Lee et al. 2015), and have less brain activation in regions related to discursive thoughts and more activation in regions related to response inhibition and attention (Brefczynski-Lewis et al. 2007).

Meditation is also associated with structural changes in the brain anatomy. Several studies have found that experienced meditators have greater gray matter concentration in specific areas of the brain. In particular, the right anterior insula, which is associated with body awareness, showed a larger concentration of gray matter in meditators (Lazar et al. 2005). Meditators also had more gray matter in the left inferior temporal gyrus and right hippocampus (Hölzel et al. 2008). In addition, meditators who practiced for longer periods of time showed greater cortical thickness in the dorsal anterior cingulate cortex, a region of the brain associated with attention (Grant et al. 2013). These results show that long-term meditation can lead to changes in specific areas of the brain that may improve attention and alleviate symptoms of ADHD disorder (Bush et al. 1999).

In terms of the effect of meditation on the autonomic nervous system activity changes, there is mixed evidence. Some studies indicate that meditation can affect autonomic nervous system activity, leading to a relaxed state with lower physiological arousal, systolic blood pressure, and heart rate (Delmonte 1984), and report the increased HRV in the low-frequency (LF) and high-frequency (HF) bands during meditation (Nesvold et al. 2012). Wu and Lo found that Zen-meditators had low, compared to the control, LF/HF ratio and LF and an increase of HF (Wu and Lo 2008). There is enough evidence that meditation can positively correlate with parasympathetic activation. However, some meditation techniques promote an active rather than a calm state. For example, in a study with ten experienced Yogi Bhajan meditators, there was a significant increase in the average heart rate relative to the baseline and a significant decrease in the coherence between heart rate and respiration (Peng et al. 2004).

A number of studies have been conducted to investigate changes in brain activity that occur during meditation. The simultaneous increase in the power of theta and alpha rhythms is considered a typical change during the practice of meditation and may indicate a state of relaxed alertness (Lomas, Ivtzan, and Fu 2015). According to Lomas’ review, mindfulness meditation has been shown in 18 studies to enhance alpha activity, and in eight studies to enhance theta activity during mindfulness meditation, in comparison with an eyes-closed resting state in experienced meditators. Only two studies reported a greater beta rhythm amplitude during mindfulness meditation. There are also several MEG studies on meditation. One study recorded data from two experienced Buddhist meditators with over 15,750 hours of practice and found changes in the anterior and posterior cingulate cortex during vipassana meditation (Calvetti et al. 2021). Another MEG study with 12 experienced participants in mindfulness meditation found that the weakening of self-awareness was associated with a decrease in gamma and beta power in certain brain areas (Dor-Ziderman et al. 2013).

There are also numerous studies dedicated to exploring the effects of meditation training on individuals just starting their meditation practice. A recent study examined the effects of 40 days of meditation training on the brain of novices (Yang et al. 2019). Meditation was associated with changes in the precuneus area of the default mode network. It was found that meditation is associated with an increased cortical thickness. A reduction in the amplitude of low-frequency waves on the EEG has been observed too. Novices participated in a six week-long mindfulness-based training and after intervention there was a significant increase in cortical thickness in the posterior left insula, an area that plays a role in auditory perception and processing (Mooneyham et al. 2017). Ten beginners underwent concentration and mindfulness meditation training. As a result, the mean alpha (central/posterior cortex) and beta 1 (cerebral cortex) amplitudes were higher in the state of mindfulness meditation than in the state of relaxation (Dunn, Hartigan, and Mikulas 1999).

Another study found two different physiological strategies for meditation in experienced meditators (Volodina et al. 2021a). The subgroup of “relaxed” meditators had higher heart rate variability coefficients—Standard Deviation of the Normal-to-Normal (SDNN) and coefficient of variation (CV), a lower stress index (SI) and respiratory rate. There was also a decrease in the frontal delta power and an increase in the occipital, parietal alpha power and alpha/theta ratio, but the overall modulation of EEG parameters during the meditation session was weak. A subgroup of “concentrated” meditators had a decrease in the heart rate and an increase in SI. During meditation, they had increasing delta power and strongly decreasing alpha power and alpha/theta and alpha/beta ratios. Both subgroups differed from beginners, but in the subgroup of the “relaxed” meditators, the difference was mainly due to peripheral indicators, and in the “concentrated” meditators, mainly due to the EEG indicators. Although all participants performed the same meditation during the experiment, their prior experiences differed in terms of what type of meditation they typically practiced.

The amount of knowledge about the physiology of meditation, both for beginners and experienced meditators, is already quite substantial. However the long term effects of meditation in novices and the amount of training required for objectively observed changes in the physiological indices to occur received little attention in the literature. Understanding the time frame associated with meditation practice is critical for planning the training activity to promote meditative practices in the modern world in order to cope with the increasing amount of stress and promote well-being.

The aim of the present study was to answer the question whether or not the minimal meditation training that can fit in the daily routine leads to the objectively observed changes in the activity of the nervous system of the practitioners both during the meditation and long term.

## 2. Methods

### 2.1. Subjects

The study initially included 38 participants who underwent pretesting, with 25 participants included in the final analysis. Thirteen participants left the experiment for personal reasons (ten people) or due to the occurrence of undesirable side effects (three people). The final sample included a 12-person group (ranging in age from 20 to 37 years, with three men and nine women and the mean age of 28.08 ± 5.45) and the 13-person control group (ranging in age from 21 to 38 years, with four men and nine women and the mean age of 27.69 ± 5.68). No participants had any prior experience with meditation, and did not have a diagnosed mental illness or brain disorder, and were not taking any drugs that affect the central nervous system (e.g. antidepressants, sedatives, etc.). Before the experiment, we analyzed various physiological parameters, as well as gender and age to ensure the meditators and the control group were similar in all indicators. The experiment was conducted in accordance with the declaration of Helsinki. Participation in the study was voluntary. All participants provided written consent approved by The HSE University Committee on Inter-University Surveys and Ethical Assessment of Empirical Research in accordance with the Declaration of Helsinki. All experimental protocols were approved by The HSE University Committee on Inter-University Surveys and Ethical Assessment of Empirical Research in accordance with the Declaration of Helsinki.

### 2.2. Experimental protocol

The design of the experiment is shown in Fig. 1. For eight weeks, participants attended 1 hour long group meetings twice a week during which they engaged in a course of Taoist meditation with an qualified instructor (experimental group) or listened to audiobooks (control group, audiobooks: “A Warm Cup on a Cold Day - How Physical Sensations Affect Our Solutions,” Talma Lobel and “Healthy Brain, Happy Life: A Personal Program to Activate Your Brain and Do Everything Better”, Wendy Suzuki). Both groups took the course/lectures in the same auditorium, in the evening for 1 hour, 2 days per a week.

**Fig. 1.**
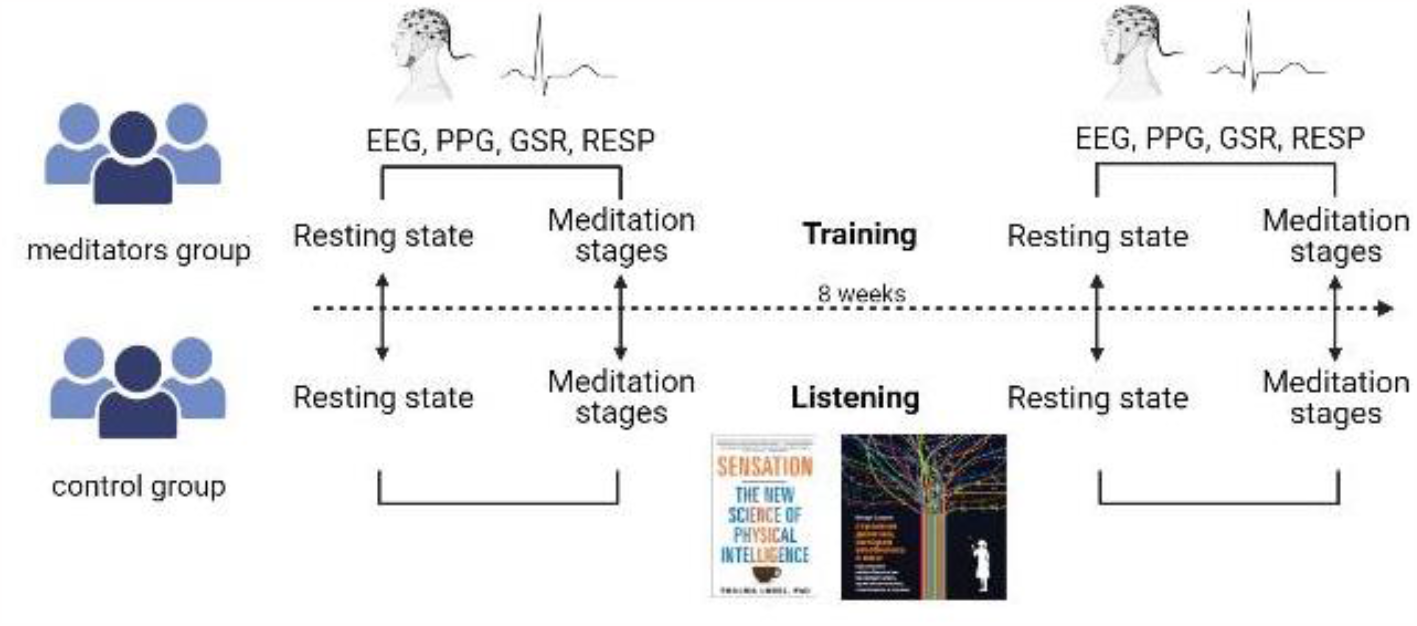
The design of the experiment with physiological testing before and after the course/listening to audiobooks with an interval of eight weeks. The first instruction was used to collect baseline resting state data, immediately after was the full meditation session ( in ∼38 min). The testing included EEG, respiration (RESP), Photoplethysmography (PPG), Galvanic Skin Response (GSR).

All the participants were required to pass physiological testing before and after the course. During such testing the experimenter placed the EEG cap, PPG, GSR, and RESP sensors and measured physiological indicators during the resting state (4 minutes: 2 minutes eyes-open + 2 minutes eyes-closed) and during the Taoist meditation guided by audio instruction, delivered through the earphones (34 minutes).

Meditation protocol during testing. Prior to the actual meditation all the participants read the text of the meditation and could ask questions. The audio instruction was presented by an experienced meditation teacher and consisted of 16 stages. The audio instruction was also used in a previous study (Volodina et al. 2021a). The end of each audio instruction marked the beginning of a new stage of meditation, and each stage lasted for about 2 minutes. For the subsequent statistical analysis we merged meditation intervals into the following 6 stages:

Resting state: Premeditation resting state;

Stage 1: Merged stages 1–4 (relaxation, body scan, taking position);

Stage 2: Merged stages 5–7 (stopping internal dialogue);

Stage 3: Merged stages 8–10 (visualization);

Stage 4: Merged stages 11–14 (coming back, focus on breathing and body);

Stage 5: Post Meditation resting stage.

The full text of meditation guidelines can be found in the Supplementary materials.

### 2.3. Physiological measurement and equipment

#### 2.3.1. EEG

The EEG data was recorded using a 30-channel wireless EEG system (SmartBCI, Mitsar, Russia) with a sampling rate of 250 Hz. The digital averaged ear signal served as a reference for all EEG data channels. The EEG data was precisely synchronized with the audio instruction. A Python script was used to simultaneously run the audio instruction and to collect EEG data.

#### 2.3.2. Photoplethysmography, Galvanic Skin Response, Respirogram

PPG, GSR, RESP signals were recorded using PolyRec (Medical computer system, Russia). The PPG sensor was placed on the subject’s index finger of the right hand. The following filters were used: 4th order 0.5 Hz high-pass filter, 4th order 10 Hz low-pass Butterworth filter, notch filter at 50 Hz. The GSR sensors were placed on the subject’s first phalanges of the ring and index fingers of the left hand. The following filters were applied to the GSR signals: 4th order 10 Hz low-pass Butterworth filter, notch filter 50 Hz.

The respirogram was measured with a thermometric sensor of nasal respiration placed under the nose whose signal was filtered in the (0.05–10) Hz range using the 4th order Butterworth filter and a notch filter centered at 50 Hz.

### 2.4. Data pre-processing

#### 2.4.1. EEG

EEG data was filtered with a bandpass filter with a lower cutoff of 1 Hz and an upper cutoff of 40 Hz. A 50 Hz notch filter was also applied to suppress the powerline interference. Ocular and muscular artifacts were removed using Independent component analysis (Urrestarazu and Iriarte 2005).

### 2.5. Data processing

#### 2.5.1. EEG

MNE-python was used for all subsequent data processing. The Welch method was applied to compute the EEG spectral power with 1 s window size without overlap. The power spectral density was then summarized by a simple averaging within the following frequency bands: Delta (1–4) Hz, Theta (4–8) Hz, Alpha (8–14) Hz, Beta (14–25) Hz, and Gamma (25-40) Hz.

#### 2.5.2. PPG

Heart rate variability indices were analyzed in the same way as in a previous article (Volodina, Smetanin, Lebedev, et al. 2021). Briefly, indices are defined as follows:

RR is the interval between successive PPG peaks, ME is the Median RR, AME is the amplitude of median, SD is the standard deviation.

1. The heart beats were detected with a commonly used peak detection algorithm implemented in routine (Camm et al. 1996). The heart rate (HR) values were calculated using a 20-second long moving sliding window with a 5-second stride. Heart rate variability (HRV) analysis is one of the commonly used methods to appraise the activity of the sympathetic and parasympathetic nervous system
2. HRV was calculated using pulse-to-pulse (RR) time series using a 60-second sliding window with the stride of 30 seconds (30 seconds overlap). Then the obtained values within each meditation stage were averaged. The HRV-based metrics included the time domain and frequency domain indices.
3. RMSSD is the “sequential difference mean squared”, the square root of the mean of the squared successive differences between neighboring NNs (beat-to-beat intervals). High RMSSD values indicate high parasympathetic activation.
4. Autonomic balance index (ABI) = AME / SD indicates the ratio between the activity of the sympathetic and parasympathetic divisions of the autonomic nervous system (Natarajan 2022).
5. Vegetative rhythm indicator (VRI) = 1 / (ME*SD) assess the vegetative balance. A lower VRI indicates greater activity in the parasympathetic nervous system, which suggests the shift towards a more balanced state of the autonomic nervous system (Moldabek 2011).
6. Stress index (SI) = AME*100% / (2*ME*dRR), which is a geometric measure of HRV that reflects the stress experienced by the cardiovascular system (Ali et al. 2021). High SI values indicate reduced variability and high sympathetic activation of the heart.

#### 2.5.3. GSR

A zero-phase high-pass 4th order Butterworth filter with a cutoff frequency of 0.05 Hz was applied to the data to remove the slow trend and the number of spontaneous reactions was measured, defined as signal fluctuations with an amplitude greater than one standard deviation of the signal calculated for each individual subject during the entire recording. A sliding window with a length of 60 seconds with 5-second strides was used.

#### 2.5.4. RESP

A peak detection algorithm was applied to the recorded breathing data. The respiratory rate was calculated using a moving window with a length of 20 seconds (strides of 5 seconds), all stages were then averaged. The respiration amplitude was calculated as the average difference between the upper and lower signal envelopes.

### 2.6. Statistical data analysis

#### 2.6.1. EEG

To determine the statistical significance of the changes in EEG power between the pre- and post-training conditions in the meditator and control group, we performed a sensor-level cluster-based permutation test (Maris and Oostenveld 2007).

To analyze changes observed during the resting state the average resting EEG power after the intervention across the frequency band for each channel was normalized to pre-intervention values. During the normalization process, the data were divided by the pre-intervention values to determine the relative change. Clusters of electrodes that exhibited significantly different EEG power between the preintervention and post-intervention states were found separately in meditator and control groups. A cluster-defining threshold of five was used. Average EEG power across the significant cluster was calculated to estimate the direction of changes.

During the analysis of EEG signal dynamics within the actual meditation process all values were standardized to the baseline measurement obtained during eyes-closed conditions. We then examined changes in EEG power across various stages of meditation in comparison to the baseline. To determine whether the dynamics during meditation were influenced by the intervention, a cluster-based permutation test was employed.

#### 2.6.2. PPG, GSR, and RESP data

Firstly we analyzed the interaction of “group” (control/meditator) and “time point” (pre-/post-training) factors using the Linear Mixed Effects model (Oberg and Mahoney 2007) for resting state data and interaction for “group”, “time point”, and “meditation stage” factors for meditation data. For indicators with a significant interaction of factors, the data before and after the intervention were compared using the Wilcoxon test with *p*-values corrected for multiple comparisons using the FDR correction procedure (Benjamini and Hochberg 1995).

## 3. Results

### 3.1. EEG

#### Resting state

We utilized a cluster-based permutation test to investigate variations in EEG power during the resting state across various frequency bands before and after the intervention. First, the post-intervention average resting EEG power across the frequency bands for each channel were normalized to the corresponding pre-intervention values to determine the relative change. Such analysis was performed for the eyes-open and eyes-closed resting state separately.

As shown in Fig. 2A our findings suggest that primarily the meditator group exhibited significant clusters, which may indicate changes in neural activity as a result of the meditation training. Specifically, our findings demonstrate that meditators showed significant clusters in the delta band in the right centroparietal area in the eyes-open condition (p = 0.01), which moved to the occipito-parietal area in the eyesclosed condition (p = 0.006). There was a big cluster comprising 22 electrodes occupying a large connected region of the scalp that demonstrated changes during the eyes-open condition (p = 0.001) that moved to the frontal and fronto-central area in eyes-closed state (p = 0.001). Alpha power analysis found another extended cluster that included electrodes in the frontal, prefrontal and left parietal areas (p = 0.001). The extent of this cluster reduced slightly in the eyes-closed condition. Beta power analysis revealed a significant cluster in the central area in open-eyes state (p = 0.032) that shifted to the left with eyes closure (p = 0.045). As barplots located next to each topographic map show, all significant clusters in meditators corresponded to the increased EEG power. In contrast, the control group only exhibited significant post-vs. pre-intervention differences in the delta band in the parietal area (p = 0.033) and the gamma band in the left prefrontal area (p = 0.032) in the eyes-closed condition (see Fig. 2B). There was a decrease in delta power and an increase in gamma power.

**Fig. 2.**
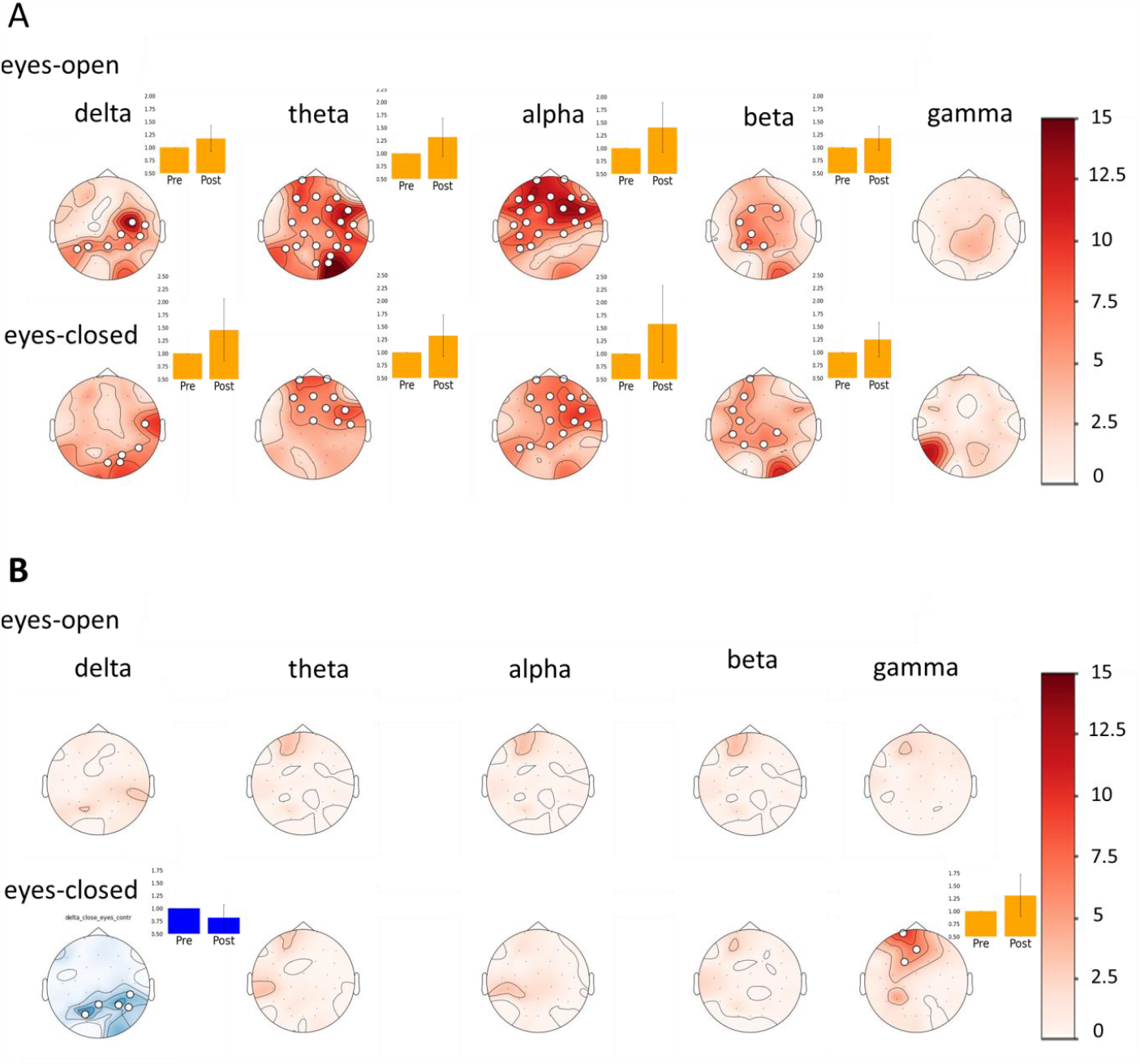
Analysis of variations in EEG power during the resting state across various frequency bands before and after the intervention using a cluster-based permutation test. The results are presented as topographic maps of the F-score. The upper panel (A) shows the results in the meditator group. The lower panel (B) shows the results in the control group. Each map corresponds to one statistically significant cluster (p < 0.05). The channels comprising each cluster are marked with white circles. For significant clusters, the barplots are presented showing the average rhythm power within the cluster±the standard deviation before and after the intervention.

#### Meditation

To conduct the initial analysis of the EEG dynamics of individuals during meditation, we compared the average pre- and post-intervention values averaged across all electrodes in separate frequency bands. Before averaging, the values of each stage were normalized to the value at rest to show the changes of EEG power during meditation. Fig. 3 shows the normalized total power profiles and there are no significant differences in the pre-vs. post-intervention curves. To further investigate the effects of the intervention on the dynamics of power changes during the actual meditation a full-fledged space-time cluster-based permutation test was conducted. However, no significant clusters were found.

**Fig. 3.**
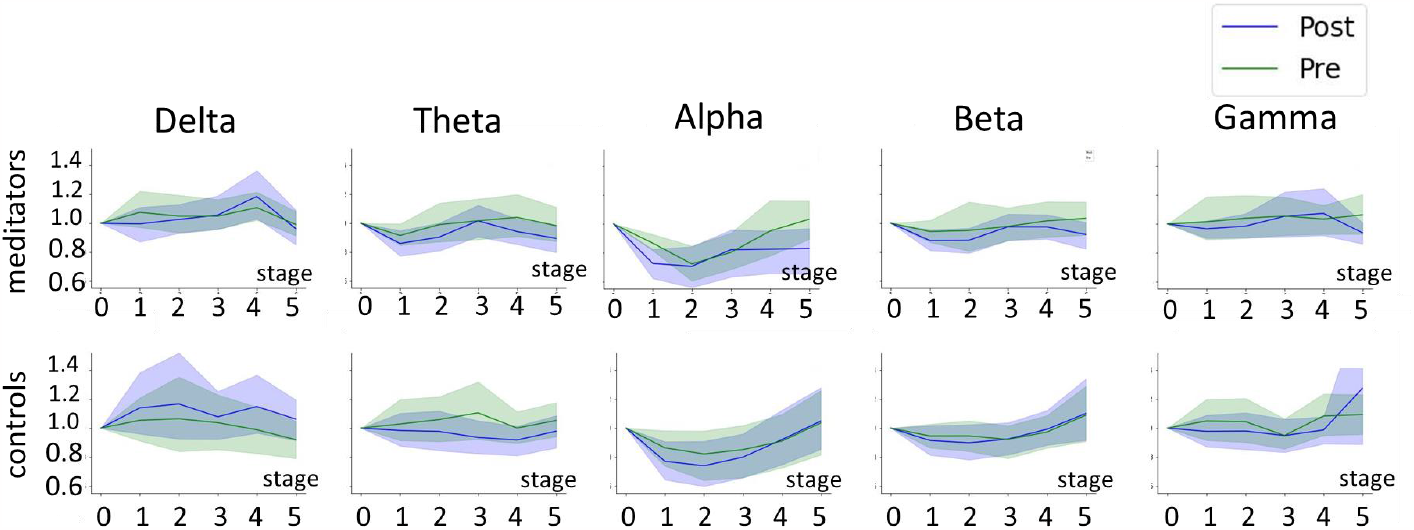
Changes in the EEG power in the process of meditation compared to eyes-closed resting state averaged across all the channels. The green line marks pre-intervention condition and the blue line marks post-intervention condition. The data are presented as the mean ± 95% confidence interval. The x-axis corresponds to the meditation stages. Stage 0 corresponds to the eyes-closed resting state condition.

Although the intervention had a clear effect on the resting state brain activity mainly in the meditator group, it did not affect the dynamics of the neurophysiological markers during the actual meditation process.

### 3.2. Autonomous nervous system markers

#### 3.2.1. Resting state with eyes-open condition

Refer to Fig. 4 here. Using the Linear Mixed Effects model we found a significant interaction of the time point and group factors for the autonomic balance index (ABI), that indicates the ratio between the sympathetic and parasympathetic activity (estimate = 0.19, z = 2.90, p = 0.03), the stress index (SI), that reflects the stress of the cardiovascular system (estimate = 0.0002, z = 3.09, p = 0.028) and the vegetative rhythm indicator (VRI), that assesses the vegetative balance (estimate = 0.007, z = 3.08, p = 0.028). See “Methods” section for the mathematical expressions used to calculate these values and the appropriate references. The post-hoc Wilcoxon signed-rank tests revealed a significant increase in ABI (p = 0.003) SI (p = 0.003) and VRI (p = 0.004), significant changes of these variables in the meditator, but not in the control group, see the corresponding error-bars in Fig. 4.

**Fig. 4.**
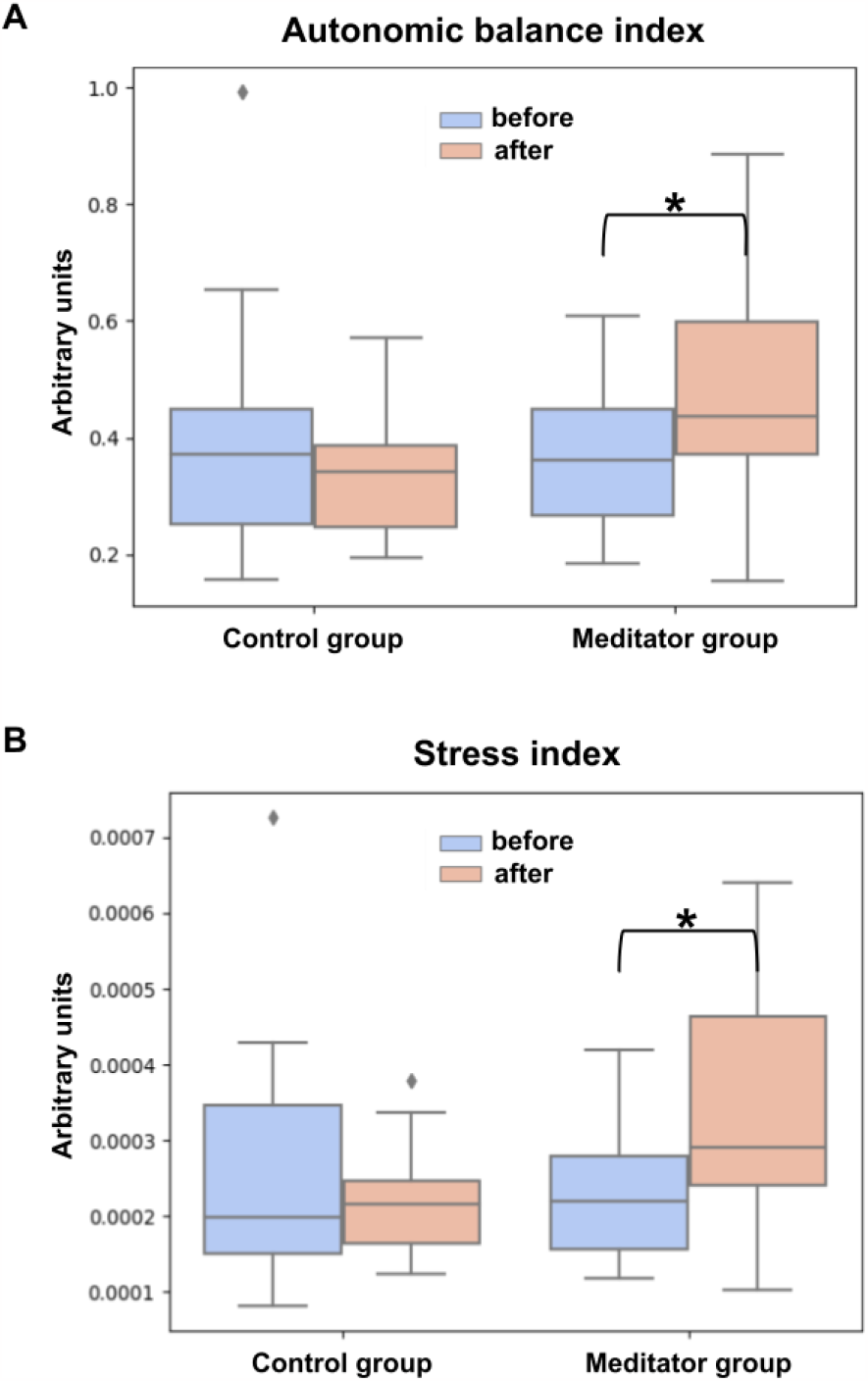
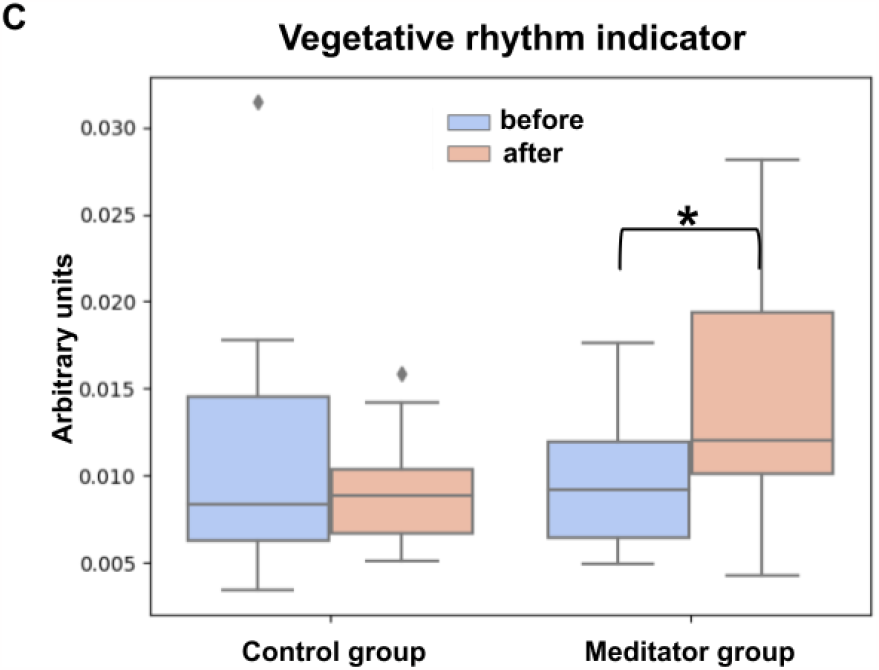
The differences between groups in PPG data during the eyes-open resting state (A is the Autonomic balance index, B is the Stress index, C is the Vegetative rhythm indicator). Data are presented as median ± interquartile range. * indicates significant (p<0.01) differences between the meditator group and the control group according to the Wilcoxon signed-rank tests followed by FDR correction.

#### 3.2.2. Resting state with eyes-closed condition

The Linear Mixed Effects model did not reveal significant interaction of time point and group factors for markers of the autonomic nervous system activity during the eyes-closed resting state in either the meditator or control group. There was no interaction of group and time point factors for respiration rate and GSR during either the eyes-open or eyes-closed resting state as well.

#### 3.2.3. Meditation

We also found a significant interaction of the time point and group factors for ABI (estimate = 0.14, z = 4.11, p < 0.001), Sl (estimate = 0.0001, z = 4.75, p < 0.001) and HR (estimate = 5.49, z = 2.77, p = 0.04). Analyzing “time point” effect separately in the groups, we found significant effect of time point only in meditator on the HR (estimate = 5.82, z = 4.31, p < 0.001), ABI (estimate = 0.12, z = 5.22, p < 0.001), SI (estimate = 9.8*e-5, z = 5.93, p < 0.001). For EEG activity we did not observe any statistically significant interaction of “time point”, “meditation stage”, or “group”. We revealed a significant interaction between “group” and “time point” factors (estimate = 1.94, z = 2.68, p = 0.01) and “group”, “time point” and “meditation stage” for the number of spontaneous galvanic skin reactions (estimate = -0.40, z = -2,01, p = 0.09). The observed interaction is brought about by the increased number of spontaneous galvanic skin reactions on the final stages of meditation in the control group after intervention. However, a more detailed post-hoc analysis using the Wilcoxon test did not reveal significant change in GSR in the control group at any stage.

There was also no effect of intervention on the respiration rate.

To summarize, we observed that after the training, the meditator group showed a trend towards an increase in EEG power across a wide frequency range during meditation, but there were no significant changes in the dynamics of the EEG and PPG indices in the process of meditation after the course. We also did not find any division of the meditator group into subgroups with different changes in the EEG and PPG indices during meditation neither before nor after the training.

## 4. Discussion

The purpose of our study was to investigate the impact of Taoist meditation training on the activity of the central nervous system (CNS) and autonomic nervous system (ANS). CNS activity was evaluated using electroencephalography (EEG), while a combination of respirometry (RESP), photoplethysmography (PPG), and electrodermal activity measurement was employed to assess ANS activity. We aimed at simulating a real-life scenario in which contemporary urban dwellers engage in meditation practices. The meditator group underwent a 16-session Taoist meditation training, each lasting an hour, over an 8-week period. The subjects were instructed not to practice meditation at home, to ensure the consistency of the intervention. The control group spent an equivalent amount of time attending offline group sessions where the participants listened to audio books. It was important for the participants to receive offline training under the guidance of an experienced instructor, who had the ability to adjust the preparatory exercises and body position. Working with an experienced instructor minimizes the risk of undesirable consequences from meditation practice (Farias, Maraldi, and Wallenkampf 2020).

As revealed by our study, Taoist meditation training had a significant effect on the baseline physiological indicators during resting state, but did not affect the meditation process itself. More specifically, the results of the study showed an increase in delta, theta, alpha and beta power during the eyes-open and eyes-closed resting state after the intervention in the meditator group but not in the control group. The results also showed changes in the heart rate variability indices associated with increased sympathetic activity and heightened alertness of the body (such as ABI, VRI, and SI) during resting state with open eyes in the meditator group after the intervention.

The increase in delta power in the meditator group observed in our study is consistent with previous research (P. L. Faber et al. 2008), which also reported an increase in delta power after meditation. Faber interpreted the increased delta activity in the medial prefrontal cortex of Zen meditators during the resting state as the inhibition of the medial prefrontal cortex, resulting in reduced emotional and cognitive engagement.

Many researchers, however, consider delta waves as a rhythm associated with cognitive processes (Knyazev 2012). Several studies have found links between delta activity and engagement in internal attention processes ((Schroeder and Lakatos 2009) (Lakatos et al. 2008) (Harmony et al. 1996)) as well as with homeostatic processes and basal metabolic rate (Boord et al., 2007, Alper et al. 2006). In particular Boord et al. (2007) found a positive correlation between basal metabolic rate and delta power recorded in 1,831 healthy subjects aged 6–86 years in the eyes open condition (Boord et al., 2007). Alper et al. revealed a positive correlation between delta EEG power and medial frontal cortical metabolism (Alper et al. 2006). There is evidence supporting the theory that delta oscillations are associated with basic motivational processes and may be involved in constant screening of the internal and external stimuli, including the stimuli that are below the threshold of conscious perception (Knyazev 2012). In our study, we observed increased delta power in the right centro-parietal area, which may be related to the activation of specific brain regions such as the insula, pre/supplementary motor cortices, and dorsal anterior cingulate cortex, as suggested by Fox (Fox et al. 2016).

Since, in our study, the increase of delta rhythm power was combined with the activation of the sympathetic nervous system, we consider such changes as signs of the activation of homeostatic processes and of brain-body interaction in meditators. Tonic sympathetic activity has been shown to support resting metabolic rate in healthy adults in previous research (Monroe et al. 2001) and increased brain-body interaction have been reported in experienced meditators in several studies (Volodina et al. 2021b), (Lutz et al. 2008).

It is worth noting that there is inconclusive data from previous studies regarding sympathetic activation as a result of meditation training. There is a common belief that meditation leads to the decreased breathing (Ahani et al. 2014) and increased parasympathetic activity (Wu and Lo 2008), (Takahashi et al. 2005), leading to relaxation. However, there is also evidence that meditation involves both the sympathetic and parasympathetic systems, with a balanced coordination of both systems needed to maintain a meditative state. A previous study revealed two different physiological strategies of meditation characterized by different physiological changes (Volodina, Smetanin, Lebedev, et al. 2021). It has also been found that different types of meditation can affect autonomic nervous system activity in different ways (Lumma, Kok, and Singer 2015), (Lang et al. 1979), (Telles et al. 2019). Tang hypothesized that different levels of meditative experience may result in different biological or physiological signatures, possibly involving the sympathetic system first and the parasympathetic system later (Tang and Tang 2020). Further research, with a longer meditation intervention, could be used to test this hypothesis.

In addition to the increase in delta power, meditators also showed an increase in theta and alpha power. Such changes observed in the meditator group after the intervention are typical changes which have been previously reported in studies of meditation (Lomas, Ivtzan, and Fu 2015), (L. Aftanas and Golosheykin 2005). One of the interpretations states that the increased alpha fluctuations reflect engagement in the internally directed tasks (Berger and Davelaar 2018). In Takahashi’s EEG based study, slow alpha power increased significantly during meditation, reflecting processes such as anticipation and attention (Takahashi et al. 2005). Theta power has been linked to attention and arousal (Başar et al. 2001), (Wascher et al. 2014), memory (Klimesch 1999), affective processing mechanisms (L. I. Aftanas et al. 1998), regulation of focused attention (Doppelmayr, Finkenzeller, and Sauseng 2008), conscious awareness (Klimesch et al. 2001), sustained attention, and mental effort (Sauseng et al. 2007). The simultaneous increase of alpha and theta power has been associated with “relaxed vigilance” (Britton et al. 2014). Studies have shown that the most reliable effects of a positive emotional state and internalized attention during meditation are reflected by increased local theta power and lower alpha power (L. I. Aftanas and Golocheikine 2001).

Moreover, the meditator group in our research also showed increased beta power, which is typically associated with sensorimotor processing (Symons et al. 2016), attention, emotions, and cognitive control (Güntekin et al. 2013). Thus an increase in beta power may indicate heightened cognitive processing and increased mental alertness. The changes in beta power were previously found during meditation in several studies. However, the direction of these changes varied. While some studies showed an increase of beta power during meditation (Thomas, Jamieson, and Cohen 2014) (Schoenberg et al. 2018), others found evidence of decreased beta power in prefrontal and parietal areas during meditation (Dor-Ziderman et al. 2013), (Pascal L. Faber et al. 2015).

Regarding the changes in the control group, we observed a decrease in delta power and an increase in gamma power during the eyes-closed resting state. We hypothesize that when listening to audiobooks, which requires attention and the activation of cognitive processes, the brain may shift into a state of heightened alertness and concentration. Individuals actively process incoming information and engage in cognitive processes unrelated to relaxation. This finding is consistent with previous research where a decrease in delta power is moderately associated with faster decision speed and higher accuracy (Van Strien et al. 2005). The observed Increase in the high-gamma frequency range is probably due to muscular activity (Muthukumaraswamy 2013).

We did not reveal significant differences in physiological measures changes during the actual meditation. The trajectory of such changes was unaffected by the intervention. Thus, we cannot make conclusions about the existence of a predisposition to certain physiological strategies of meditation (Volodina et al. 2021a).

We can conclude that 16 hours of instructed meditation training over 8 weeks is not enough to get a stable physiological effect on the meditation process itself. However, even this relatively short intervention resulted in changes in the activity of the central and autonomic nervous system. These changes may indicate improved attention, particularly attention to internal cues, and increased alertness.

Nevertheless, the results of the study indicate that a longer or more consistent practice of meditation may be required within the experiment in order to form a solid foundation for creating an effective device for assisted meditation training. This suggests that further research with longer intervention periods is needed to better understand the physiological effects of meditation and develop more effective meditation training methods.

## 5. Conclusion

The course of Taoist meditation, consisting of 16 hours of training over 8 weeks, caused changes in brain activity (increase in delta, theta, alpha, beta power) and markers of autonomic nervous system activity which indicate sympathetic activation. The effect was primarily observed in the resting state. There were no significant changes in the dynamics of indicators during the actual meditation, which could be due to the insufficient duration of the training. The trajectory of physiological changes during the meditation did not change after the course. Since we did not observe changes in the physiological indicators during the actual meditation process, we could not draw conclusions about the existence of a predisposition to one of the meditation strategies. Further research is needed to explore this question.

As shown in Fig.3 during the actual meditation process, we could observe clear stage-by-stage variations in the band power which forms frequency-specific oscillatory power profiles. We found no changes between these profiles for the experimental and control groups. This conceptually complicates the development of assistive devices aimed at “guiding” novice meditators during the actual meditation process. According to our results, the focus in creating such digital assistants should be shifted towards monitoring neurophysiological activity during the time intervals outside of the actual meditation. As apparent from Fig. 4 these changes occur not only in the EEG derived parameters but are also detectable based on the markers of autonomous nervous system activity which can be readily measured with a range of wearable devices which renders hope for a rapid translation of our results into practical applications.

## Availability of data

The data used in this study is available upon request from the corresponding author—Maria Volodina.

## Acknowledgements

The research leading to these results has received funding from the Basic Research Program at the National Research University Higher School of Economics.

## Author contributions statement

Maria Volodina conceptualized and designed the experiment, Anna Rusinova and Maria Volodina performed the experiment, analyzed the data, and wrote the article. Alexey Ossadtchi conceptualized and supervised data analysis, edited the article and acquired funds. All authors reviewed the manuscript.

## Additional information

Correspondence and requests for materials should be addressed to Maria Volodina.

The corresponding author is responsible for submitting a competing interests statement on behalf of all authors of the paper.

## Data and code availability statement

Raw EEG, PPG, GSR, RESP data, supplementary tables and python code will be publicly available in the Open Science Framework webpage.

See https://osf.io/3y4nj/?view_only=a429a88daf7a48f8af58f9c767e37dc6.

