## Supplementary materials for "Short-Term Meditation Training Alters Brain Activity and Sympathetic Responses at Rest, but not during the meditation"

Full text of the meditation guideline.

Resting state for 2 minutes with open eyes:

“Let's begin the experiment. Get into a comfortable position. Sit with your eyes open.”

Resting state for 2 minutes with close eyes:

“You should take a comfortable position and close your eyes.”

Stage 1 (2 min):

“The beginning of meditation. Sit up straight. Straighten your back. The top of the head stretches to the sky, and the sacrum stretches down. Relax your facial muscles. Relax the forehead, temples, jaws, chin, neck muscles, collarbones. Shoulders, elbows flow down. Relax your ribs. With each breath they become softer. Relax your stomach and lower back. Breathing gets deeper.”

Stage 2 (2 min):

“Shift your attention to the sacrum. From it, slowly raise your attention upward, increasing the distance between the vertebrae. Sliding up: 5th lumbar vertebra, 4th, 3rd, 2nd, 1st. Stretch the thoracic spine: 12th, 11th, 10th, 9th, 8th, 7th, 6th, 5th, 4th, 3rd, 2nd, 1st. We relax the muscles along the spine, the tendons that hold it. Slide, grow up. Shoulders, elbows are relaxed, as if flowing down. The ribs become soft. Stretch your neck, also vertebrae by vertebrae. Slowly, but persistently. Shift your attention, raise your body. 7th cervical vertebra, 6th, 5th, 4th, 3rd, 2nd, 1st. Maintain this state.”

Stage 3 (2 min):

“Imagine that you are hanging from the top of your head. Pull yourself up with all your might. Even stronger and stronger. You're hanging like an empty bell. The spine is stretched. The sacrum stretches downwards, stretching the spine with its weight. The spine begins to stretch between the head and the vertebrae of the sacrum. Maintain this state.”

Stage 4 (2 min):

“The tongue touches the palate. The energy flow forms a platform at the level of the third eye. This platform is becoming more powerful, and its gravitational force is growing. Gently transfer the mind to the platform and fix it on it. Maintain this state.”

Stage 5 (2 min):

“Relax your mind and allow it to leave your body, filling the entire space of the room.”

Stage 6 (1 min):

“Direct your attention to the boundaries of your body.”

Stage 7 (2 min):

“Look inside yourself. Let your mind become aware of the inner emptiness. Relax your mind in it. Maintain this state.”

Stage 8 (1 min):

“Say to yourself: “The sky above me is open and endless.” Look at the sky and realize its infinity.”

Stage 9 (1 min):

“Say to yourself: ‘The earth is solid, powerful. Realize the hardness and power of the earth.”

Stage 10 (4 min):

“Say to yourself: ‘I am between heaven and earth. Like a pillar supporting the heavens, filling the space between heaven and earth. Powerful.’ Understand yourself as being powerful by sitting on the ground and supporting the heavens. Maintain this state.”

Stage 11 (2 min):

“Feel your body, “Qi” and “Shen” becomes infinite.”

Stage 12 (1 min):

“Realize the boundaries of your body.”

Stage 13 (1 minute)

“Direct your attention between the eyebrows, to the tip of the nose, to the center of the chest, to the solar plexus and to the lower abdomen. Starting from the surface of the abdomen, direct your attention inward. Breathe naturally.”

Stage 14 (2 min):

“Focus your attention on your stomach. Pull in the stomach on the inhale and expand on the exhale.”

Stage 15 (1 min) (state of rest after meditation):

“Breathe normally. Expand your stomach on the inhale and pull it in on the exhale.”

“When you are ready, open your eyes and finish the meditation.”
